# Short-term memory, attentional control and brain size in primates

**DOI:** 10.1101/2024.02.26.582100

**Authors:** Carel P. van Schaik, Ivo Jacobs, Judith M. Burkart, Gabriela-Alina Sauciuc, Caroline Schuppli, Tomas Persson, Zitan Song

## Abstract

Brain size variability in primates has been attributed to various domain-specific socio-ecological factors. A recently published large-scale study of short-term memory abilities in 41 primate species [1] did not find any correlations with 11 different proxies of external cognitive demands. Here we found that the interspecific variation in test performance shows correlated evolution with total brain size, with the relationship becoming tighter as species with small sample sizes were successively removed, whereas it was not predicted by the often-used encephalization quotient (EQ). In a subsample, we also found that the sizes of brain region thought to be involved in short-term memory did not predict performance better than did overall brain size. The dependence on brain size suggests that domain-general cognitive processes underlie short-term memory as tested in [1]. These results support the emerging notion that comparative studies of brain size do not generally identify domain-specific cognitive adaptations, but rather reveal varying selection on domain-general cognitive abilities. Finally, because attentional processes beyond short- term memory also affected test performance, we suggest that the delayed response test can be refined.

## 1. Introduction

For nearly half a century broad comparative studies have examined vertebrate brain size as a function of a range of variables used as proxies for likely selective pressure on the evolution of cognitive abilities. The challenges causing these selective pressures are often categorized as either social, sexual, or ecological (or more specifically navigational or technical [2]. For instance, in various vertebrate taxa, enlarged brains have been linked to pair-bond strength [3], cooperative breeding [4], female mate choice [5], social deception [6], group size [7], diet quality [7–9], vocal learning [10], home range size [11], seasonality [12], habitat complexity [13], extractive foraging [14], and technical innovations [15]. Each of these studies found significant correlations between the specific putative selective pressure and absolute brain size (or correlated neuroanatomical measures like neocortex size or neocortex ratio).

By looking for a specific cognitive challenge as the selective agent, these studies assume that cognition is domain-specific, i.e., that specific cognitive abilities arose as adaptations in response to specific cognition-based selective pressures, such as remembering where cached food is stored or assessing the suitability of potential allies in agonistic contests (cf. [16]). But such specific cognitive adaptations may need only small neural networks [17], so may not affect overall brain size. Indeed, few studies have revealed clear correlations between a specific cognitive adaptation and the size of particular brain regions (apart from the well-documented correlation between hippocampus size and spatial memory in food-caching and brood-parasitic birds [2,18,19]). If, alternatively, they do assume that these cognitive adaptations are (or have over time become) domain-general, i.e., being part of cognitive processes that are decontextualized and usefully deployed in a variety of contexts [20], it is very unlikely that they would still be linked to a specific selective cognitive demand, but rather to a great number of them. Either interpretation may therefore not be fully compelling.

This raises the question how often these comparative studies correctly identified the selective agent underlying specific cognitive adaptations. Indeed, recent studies that used larger datasets and tested multiple hypotheses (to disambiguate inter-correlated variables) found either little support for the link between overall brain size and specific social or ecological predictors [21,22], or found that multiple predictors matter, which is more suggestive of domain-general processes [7,23]. Moreover, across studies, the results are unstable when data are analysed differently [7,22,24,25], which points in the same direction. All this therefore suggests that specific selective agents rarely affect over all brain size (although they may nonetheless affect the size of specific brain regions).

By contrast, the brain’s overall size or its total number of neurons in primates and birds is a good predictor of interspecific variation in domain-general cognitive ability extracted from tasks in the lab [10,26–28] or cognitive performance and behavioural flexibility in nature [12,29–32]. Likewise, domain-general cognitive abilities are good predictors of interspecific variation in executive functions, such as inhibition in primates [20] and of quantity estimation in birds and mammals [33]). These domain-general cognitive abilities are not directly linked to any specific eco- or socio-cognitive challenge (although this does not imply that domain-specific and domain-general cognitive adaptations exclude each other). This second group of comparative studies therefore appears to have identified variation in the conditions favouring overall behavioural flexibility (reviewed in [20]).

The relative importance of domain-specific and domain-general cognitive adaptations in setting brain size is difficult to resolve in more complex species like vertebrates. To assess whether domain-specific cognitive adaptations affect total brain size, we may need very large data sets owing to large individual variability and sometimes low replicability in cognitive tests (e.g. [1,10,34,35]) and the use of proxy measures, whose correlation with the actual putative demands may be far from perfect and which may also be correlated with other demands. However, there is another, more immediate prediction: we should find that across species, estimates of the cognitive demand or challenge should predict performance on tasks estimating the domain-specific cognitive ability. If this is found, we can then look for links between either of these two and the size of specific brain regions thought to be involved in regulating the ability. If, however, no correlation between demand and performance is found, we may expect the performance to be correlated with the size of the total brain rather than species regions. If the latter is found, it suggests that the ability is part of a broader, coevolved bundle of abilities, generally expressed in domain-general cognitive abilities, such as executive functions.

A recently published study of short-term memory (STM) using the delayed-response paradigm [1] provides a fresh opportunity to examine both domain-specific and domain-general interpretations simultaneously. ManyPrimates et al. [1] tested an unprecedented 421 individuals across 41 primate species on a delayed-response task, where subjects had to indicate which cup, among three identical ones, covered a reward hidden by the experimenter 0, 15, or 30 seconds earlier. They found no correlation between performance at the three time intervals or their average value and 11 socioecological variables that were considered to represent (i.e., to be “proxies” of) cognitive demands in nature: colour vision type, day journey length, diet diversity, dietary breadth, relative feeding time, group size, home range size, percentage frugivory, diurnal resting time, terrestriality, and vocal repertoire size. These results therefore argue against the domain-specific interpretation. However, although ManyPrimates et al. [1] found a clear signature of phylogeny in performance, they did not consider the correlations with the size of the total brain, which would test the domain-general interpretation, or with the size of specific brain regions, which would test the domain-specific interpretation.

The aim of this paper is to perform this latter analysis using published neuroanatomical information. First, we examined the link between STM performance and absolute brain size and Jerison’s [36] encephalization quotient. We reasoned that if we do find a clear brain size effect on test performance (cf. [27]), this argues for STM being an expression of domain-general cognitive abilities, rather than a domain-specific cognitive adaptation to a specific ecological (e.g., remembering the presence of temporarily invisible food sources) or social (e.g., remembering the presence of a nearby but invisible rival) challenges.

Second, for a subset of species (constrained by data availability), we also examined whether STM performance could be linked to the size of specific brain regions proposed as putative neuroanatomical substrates of STM. to check. The brain regions included in these analyses were the neocortex, cerebellum, prefrontal cortex (PFC), and primary visual cortex (V1). The neocortex is traditionally considered as the seat of recently evolved complex cognition, with its notable expansion in humans driving our supposedly unique abilities [37]. Thus, neocortex size represents an additional relevant neuroanatomic variable for testing the domain-generality of STM. According to the dominant theory in the field of STM processing, persistent activity in the PFC is both *necessary* and *sufficient* to implement STM in general, with activations in the dorsolateral prefrontal cortex (DLPFC) being specific to visuospatial STM (e.g., [38]). More recent accounts, however, highlight the crucial implication in STM processing of more posterior neocortical regions (PPC, V1), as well as extra-cerebral structures (the cerebellum). The cerebellum is densely packed with neurons and expanded disproportionately early in the evolution of apes [39]. Among its many functions is visuospatial working memory [40]. The PPC has been found to have a role in both short- and long-term visuospatial memory [43, 38], while V1 grey matter volume has been found to be predictive of STM performance in humans [41]. In these recent accounts, the PFC only exerts supplementary functions in STM processing related to executive control, thereby enabling individuals to withstand interference or mere time-based decay and, thus, to hold items in memory for longer periods of time [41–43]. Whereas the PFC, the PPC and the cerebellum exhibit ‘multi-demand’ (e.g., domain-general) activation patterns, (40, 45, 46), the V1 is a more specialized sensory-processing area. Assessing the relationship between V1 size and STM performance, thus, also offers a potential avenue to explore whether a domain-specific signature of STM could be traced in primates.

Finally, we examined whether the test estimated short-term memory only, or also attention management.

## 2. Methods

We aimed to test for an effect of brain measures on the performance on the short-term memory task, described in detail in ManyPrimates et al. [1]. Since the design of the original study was uniform across species, and its analyses had already controlled, and found no evidence for, confounding effects, our statistical model could be very simple. Because females are the ecological sex, we took female brain size and female body size from Isler et al. [44], supplemented with data from [45] and [46]. Due to missing data, we took the brain mass of *Trachypithecus phayrei* to stand for that of the similar-sized *T. francoisi*, for which no brain data are available.

Effects of neuroanatomical variables were examined at two levels. First, we conducted analyses involving the following global measures: ln-transformed absolute brain mass (after correcting endocranial volume data into mass, where needed, by multiplying it by 1.036, as per Isler et al. [44], and encephalization quotient or EQ, i.e., the ratio of observed to expected brain size, based on an empirical, mammal-wide regression of log_10_ brain size on log_10_ body size [36]. EQs were estimated using Jerison’s [36] equation for expected brain size of 0.12 x P^0.667^, where P is body mass in gram. We did not consider additional global estimates, such as estimated neuron counts or estimates of the size of the cognitive brain, because these are all very highly correlated with overall brain size (r > 0.98) and given the variability in cognitive measures would not identify meaningful differences (see [50]).

Second, for a smaller subset of species due to limited data availability, we also examined the fit with an additional global measure, i.e., the neocortex, as well as three brain regions thought to be involved in STM (see introduction), namely the size of the cerebellum, and, based on the findings of [44], the amount of grey matter volume in the prefrontal cortex (PFC) and in the primary visual cortex (V1). For these analyses, we included data from different sources, but applied a priority rule based on species numbers. For cerebellum, we used Heldstab et al. [45], supplemented by DeCasien & Higham [47], and for neocortex, we used the same sources with the same rule, supplemented by Barton & Venditti [39]. For grey matter in V1, we used DeCasien & Higham [47], and for PFC we used Schoenemann et al. [48]. We then compared the statistical fit of these regions with that of the total brain. (In all these cases, the results did not depend on which source of total brain mass we used, which differed slightly among sources. However, for the analysis of overall brain size we relied on female brains only and report it also for the brain region analyses.)

The response variable was the performance on the STM test performance. Because performance declined with increasing delay [1], we took the average performance (proportion of correct choices) over the three intervals (0, 15 and 30 seconds) for the first analyses (results were substantially similar if we used the three direct measures; see Table S2).

To examine the impact of the various brain measures on the short-term memory (STM) performance, following ManyPrimates et al. [1] we used the Markov chain Monte Carlo (MCMC) generalized linear mixed model procedure, implemented in the ‘MCMCglmm’ package [49] in R [50] v. 4.2.1. We used the ‘ele_1307_sm_sa1’ phylogenetic tree from Fritz et al. [51], into which we inserted *Eulemur flavifrons* based on Meyer et al. [52]. The full dataset, consisting of 41 species, was utilized for the analyses involving total brains size and EQ. We used the priors (list(R = list(V = 1, nu = 0.002), G = list(G1 = list(V = 1, nu = 1, alpha.mu = 0, alpha.V = 1000)))) and ran the MCMC algorithm for 75,000 iterations, with thinning of 40 and burn-in of 7500. Significant counts referred to the presence of significant p-values in 100 iterations. To test the model fit in relation to the number of subjects per species we compared the R^2^ of PGLS analyses on the same data set, using ‘phylolm’ package [53].

## 3. Results

In a first test, we used the average for each species of the mean performance on the delay of 0, 15 or 30 seconds, as the response variable. The MCMCglm showed a significant statistical effect of ln brain size on performance in the STM test, but no effect of EQ (Table 1.a; Figure 1.a). However, there was a clear effect of the number of subjects tested per species on this STM estimate. The fit of the model for overall brain size, as indexed by the R^2^, improved at each step when we consecutively removed species with 1 subject, then those with 2 subjects, etc., until species averages were based on 10 or more subjects (Figure 2). Table 1.b and Figure 1.b show the results for the 11 remaining species that all had at least 10 subjects. The effect of ln brain size had become highly significant, but that of the EQ remained non-significant.

**Table 1.**
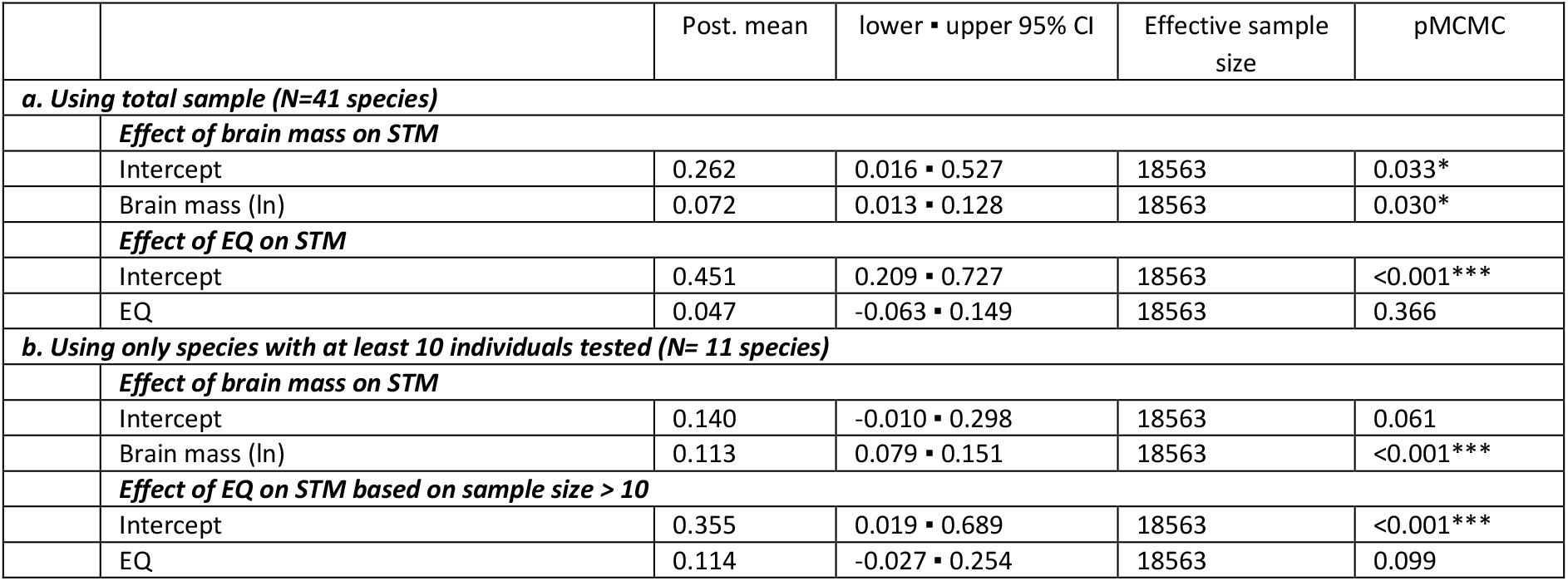
Results of the MCMCglmm for the effects of brain size or encephalization quotient (EQ) on average performance on the short-term memory test data of ManyPrimates et al. [1], for the full sample (**a**), and the reduced sample in which only species with at least 10 individuals tested were included (**b**).

**Figure 1.**
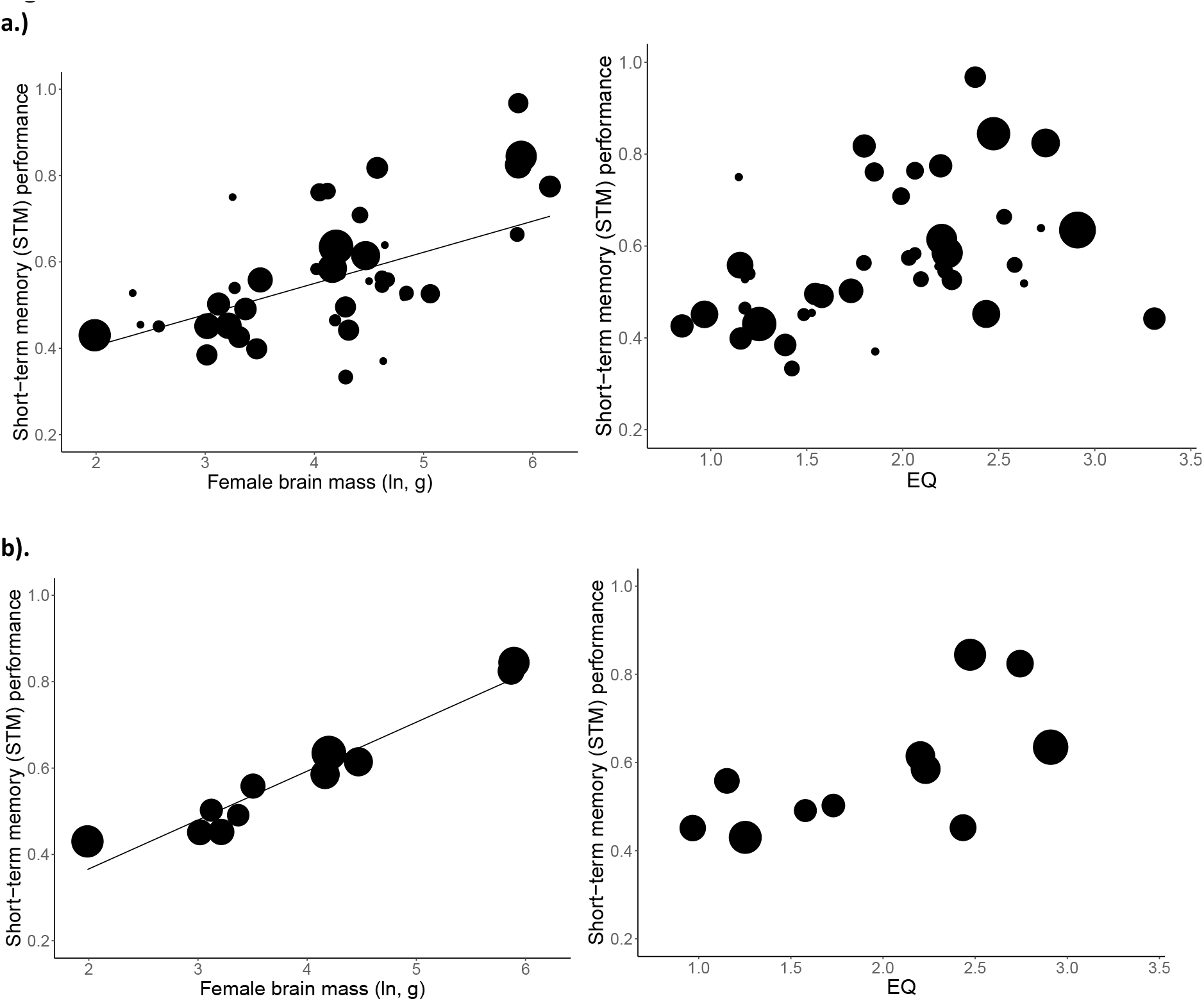
The relationship between a species’ average performance in the short-term memory test and (ln) brain size and the encephalization quotient, for the full sample (**a**), and the reduced sample in which each species had at least 10 tested subjects (**b**). Shown is a linear regression. The line indicates the relationship is significant (see Table 1). The shaded areas delimit the 95% confidence limits generated from posterior samples of the MCMCglmm. Size of the points reflects the number of subjects tested for the species concerned.

**Figure 2.**
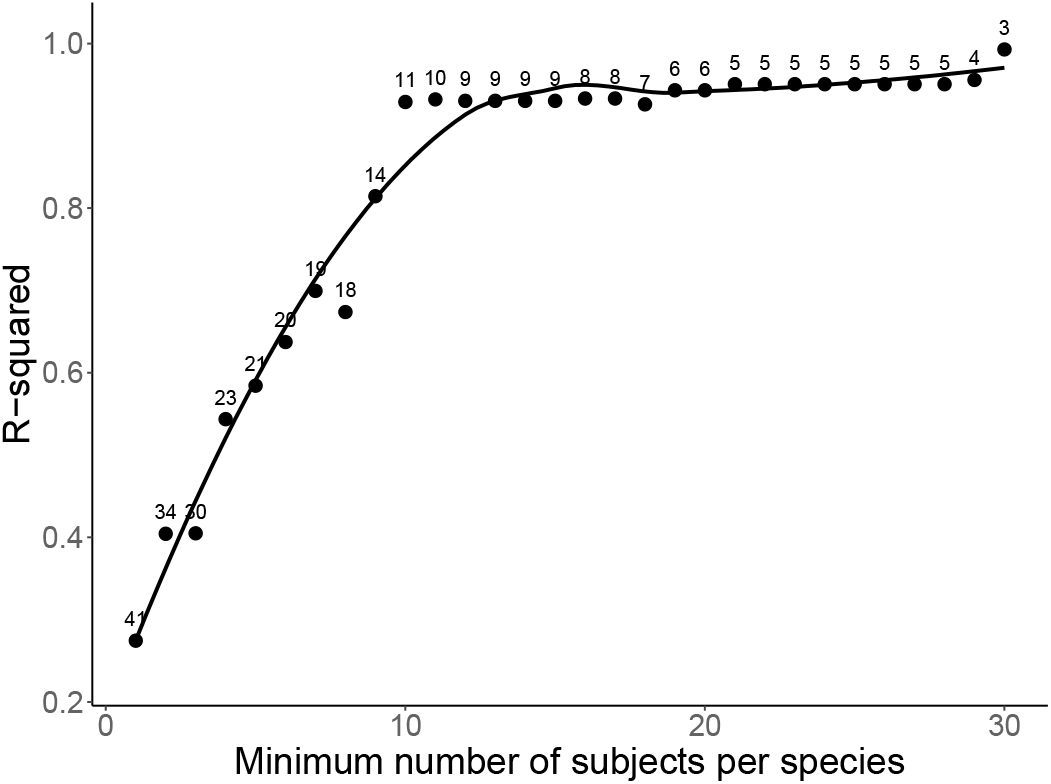
The proportion of variance in average performance on the short-term memory test explained by ln brain size in the PGLS model, indexed by R^2^ as we increase the minimally required number of subjects sampled per species. Small numbers near the points refer to the number of species in the analysis. Notice that at 10 subjects per species the fit between brain size and performance becomes near-perfect.

We next examined whether the size of specific brain regions known to be involved in memory functions were better predictors of average STM performance than total brain size. As shown in Table S1, for neocortex size the results were virtually identical as for total brain size and in the reduced sample of 11 species with larger number of subjects this remained true as well (compare with Table 1; Figure S1.a). For cerebellum, the fit with STM performance was slightly better, but this was no longer the case in the reduced sample (See Table S1, Figure S1.b). For the grey matter in the visual cortex and the prefrontal cortex, the number of species included in the analysis was smaller, which precluded formal tests.

However, in both cases total brain size tended to be the better predictor, rather than worse, which is inconsistent with the hypothesis that STM is a domain-specific ability dependent on the size of the regulating regions.

Finally, we examined the effect of the delay on performance. A delay of 0 seconds presumably imposes only modest demands on memory. We therefore expect the performance of all species to start high regardless of brain size. Figure 3.a shows this expectation when the results are fully explained by short-term memory, which in turn is (as the first set of results suggests) explained by brain size. However, this expectation was not met. In the dataset with more subjects per species (N=11), brain size already positively affects performance at the zero-seconds delay, when memory should hardly play a role (Table S2). At the same time, there was a significant effect of delay (as shown in the ManyPrimates et al. [1] study, here replicated in Table S3). However, there was no interaction between the statistical effects of overall brain size and delay (Table S4), indicating that the decline in performance with increasing delay was not affected by brain size (as illustrated in Fig. 3.b: the lines are roughly parallel).

**Figure 3.**
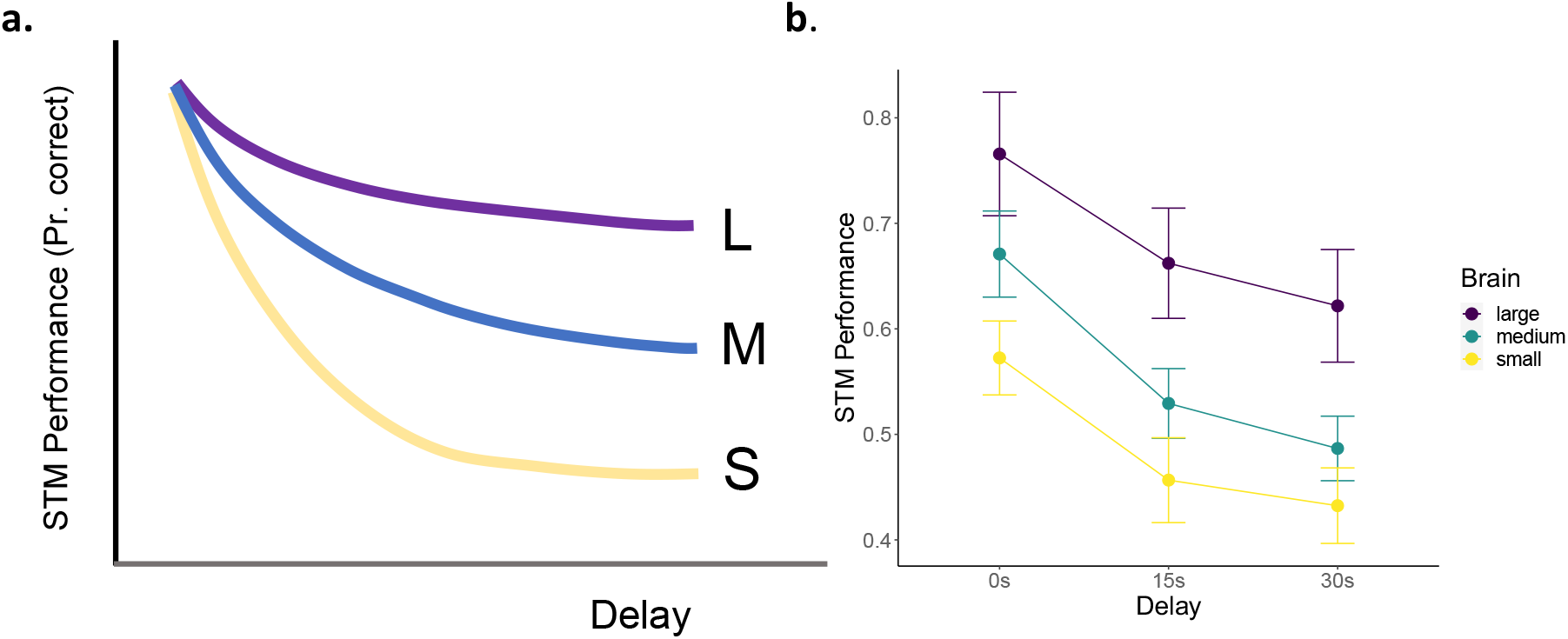
**(a**) Sketch of expected proportion correct choices (out of 3 options) in three delay categories for three classes of brain size; (**b**) observed values (in three distinct classes of brain size, which correspond to brain sizes that are greater than the 75^th^ percentile, between the 25 ^th^ and 75 ^th^ percentiles, and less than the 25 ^th^ percentile of the dataset, respectively). The vertical bars refer to ± 1 standard error of the mean. The results in (b) are merely illustrative; for statistics see model result (Table 2).

## 4. Discussion

A recently published, detailed comparative experimental study in captivity of visuo-spatial short-term memory (STM) ability in 41 species of nonhuman primates [1] did not find correlations with any of the 11 potential selective agents considered, and therefore did not find evidence for STM as a domain-specific cognitive adaptation. However, ManyPrimates et al. [1] did not pursue additional analyses to examine the possibility that STM task performance was an expression of domain-general cognitive abilities, which would suggest a link with internal predictors like brain size. Our goal here was to do these analyses.

The follow-up analysis showed that the effect of brain size on STM estimates was significant (Figure 1), as did previous work that examined performance on a wide range of cognitive tasks [27,29]. However, because the STM measure was quite variable, as is often seen in cognitive tests (e.g., [34,35]), we also controlled for the effect of the number of subjects sampled per species. The correlation between brain size and average STM performance became stronger the more species with small sample sizes were removed (Figure 2). (The ManyPrimates [1] study did not notice this effect because it had controlled for different sample sizes per species by repeatedly sampling a randomly selected single subject for any given species, based on the reasonable assumption that each individual is representative of its species.)

This sample size effect is important because it shows that the poorer predictive value of brain size in the full sample was due to individual differences within species (cf. [54]) rather than the presence or absence of specific cognitive adaptations affecting task performance in particular species. The latter case would have suggested an effect of some external selective agent. The extremely tight link with absolute brain size also made it unlikely that the statistical effect of specific brain regions on average STM performance of specific brain regions could be stronger. The additional analyses of brain region effects confirmed this, and the V1-centred analysis also provides an additional argument against a role for specific selective agents.

At present, we see three possible explanations for the finding that STM performance was tightly correlated with overall brain size but not with any of the possible selective pressures considered in the original study nor showed a better fit with the size of specific brain regions than with total brain size. First, previous work may simply have not looked hard enough for these domain-specific cognitive adaptations. However, studies examining how specific selective pressures favoured specific cognitive adaptations that affected the size of specific brain regions remain rare [55]; the best-documented cases remain the food caching and brood parasitism in birds affecting spatial memory and the relative size of the hippocampus [2,18,19]. While the present study replicated the negative results of [1] for a broad range of socioecological variables, it is still possible that the correct selective agent remains unidentified. In addition, the preliminary analyses of the link with specific brain regions, though negative, had only limited statistical power. Thus, while there is no support for this possibility, we cannot at present reject it.

Second, the effects of domain-general abilities and domain-specific cognitive adaptations on brain size may coexist, but when we checked the statistical effect on STM performance of each of the possible selective agents, there was no evidence for this, not even in the reduced set with the poorly sampled species removed (see Table S5). If the effect of domain-general cognitive abilities dwarfs that of the domain-specific ones, broad comparative analyses involving total brain size are not the right tool to examine the latter’s importance (cf. [55]. Instead, more detailed correlations between the selective agent, performance on relevant cognitive tests, and specific brain regions must be explored.

Third, seemingly modular abilities may also arise developmentally based on the application of domain-general processes in specific contexts [20]. Thus, the specific cognitive adaptations may consist of innate predispositions to attend to specific stimuli or respond in particular ways, which then during development recruit domain-general mechanisms to produce processes that are functionally similar to domain-specific cognitive adaptations (e.g., fast and seemingly informationally encapsulated).

It would be premature to draw a firm conclusion as to which of these three possibilities applies or apply in the case of the STM performance. However, the results are most consistent with the idea that general behavioural flexibility is responsible for STM performance and so explains its link with overall brain size.

Two more methodological issues also warrant discussion. First, the encephalization quotient (EQ) did not predict performance on the STM test, echoing tests of other cognitive measures [27,29,34]. The EQ is based on an empirical regression of brain size on body size [36]. However, intraspecific regressions suggest that this empirical inter-specific regression greatly overestimates the portion of the brain required to support purely somatic processes linked to body size [56], and by implication underestimates the elaboration of sensorimotor functions and their integration as well as cognition related to behavioural flexibility. This confirmation of the poor predictive value of the EQ implies we should avoid using it as an estimate of cognitive abilities (although it may work as such in ectotherms: Triki et al. [57]).

Second, the results raised the question what exactly is tested by the delayed response task. Performance declined with increasing delay, and did so non-linearly in the way expected by the Ebbinghaus effect (the “forgetting curve”: Schubiger et al. [58]), indicating that the test did estimate STM. However, brain size did not affect the steepness of this delay, and thus did not affect STM (Table S4). This is somewhat surprising, given that in humans cortical structures are strongly involved in short-term memory [38,41–43]. This pattern suggests the construct validity of the test is imperfect. Indeed, the naïve expectation (Fig. 3.a) that all species should do equally well on the delay of 0 seconds was not met. However, there may be reasons to doubt that all species experienced the same zero-second delay. First, the interval between hiding the reward and allowing the subject to choose is more than 0 seconds, albeit presumably the same for each species. Second, the latency between the opportunity to choose and the latency until the actual choice is not reported, but if smaller-brained species respond more slowly, they may in effect experience a longer delay than the nominal one. It is currently unknown how important this hidden variation in delay durations is. However, if it is not important, the brain size effect on performance may reflect the strategic deployment of attention where this may bring a food reward (cf. [1]), which can be seen as an expression of executive, top-down control (cf. [20,33,34,59–62]. That attention is a scarce resource for smaller-brained species is suggested by the result of [58] in the same test. Performance greatly improved when animals had to remember in which of 9 boxes food was hidden, instead of the far easier setup with 2 boxes. Apparently, when only 2 boxes were present, the subjects did not bother to attend in detail, suggesting that other demands on their attention prevailed.

The pattern in the results therefore suggests that the paradigm tested short-term memory as well as executive functions, presumably the strategic deployment of attention, with the latter perhaps strengthening the link with brain size. In fact, the crucial role of attention was already noted in the first study of delayed reactions over a century ago [63]. Thus, further refinement of the test format may improve construct validity (cf. [58]). It therefore cannot be excluded that future tests may show how STM performance depends on some specific environmental demand.

In sum, we found that the performance on the STM tests across a broad array of primate species was strongly predicted by brain size, whereas this was not the case for a large array of often-used proxies for specific cognitive demands tested by ManyPrimates et al. [1]. Neither did a more limited number of specific brain regions improve the model fit over that of the total brain. The results therefore support the notion that the strength of domain-general abilities, such as executive control and perhaps general intelligence, explain most of the variation in STM performance. This may be a more general pattern among primates, and perhaps other mammals and birds.

## Acknowledgments

We could not have done this study without the enormous effort of the ManyPrimates consortium. We are therefore very grateful for their generosity. The Swedish Research Council has funded the contributions to this study by Ivo Jacobs (grant 2019-03176), Tomas Persson and Gabriela-Alina Sauciuc (grant 2019-03104).

