## Supplemental tables and figure for "Short-term memory, attentional control and brain size in primates"

### Table S1.

Results of MCMCglmm analyses for the effects of grey matter size in the primary visual cortex (GM) and the size of the prefrontal cortex (PFC) on average STM performance. Note the limited samples sizes. We also repeated the analyses for neocortex and cerebellum for the reduced data set including only species with at least 10 subjects tested.

| | | Post. mean | lower ▪ upper 95% CI | Effective sample size | pMCMC |
|---|---|---|---|---|---|
| ***a. GM in V1 volume (N=17 species)*** | | | | | |
| | ***Effect of gm V1 on STM*** | | | | |
| | Intercept | 0.477 | 0.301 ▪ 0.659 | 18411 | <0.001 |
| | *V1.gm.vol.3.* (ln) | 0.076 | -0.011 ▪ 0.154 | 18563 | 0.085 |
| | ***Effect of gm V1 volume on Brain mass*** | | | | |
| | Intercept | 2.793 | 2.178 ▪ 3.424 | 18563 | <0.001*** |
| | *V1.gm.vol.3.* (ln) | 1.186 | 0.913 ▪ 1.457 | 17819 | <0.001*** |
| | ***Effect of brain mass on STM in same subsample*** | | | | |
| | Intercept | 0.262 | 0.024 ▪ 0.509 | 18563 | 0.028* |
| | *Brain* (ln) | 0.073 | 0.016 ▪ 0.123 | 18563 | 0.022* |
| ***b. GM in PFC volume (N=8 species)*** | | | | | |
| | ***Effect of gm PFC on STM*** | | | | |
| | Intercept | 0.498 | 0.190 ▪ 0.810 | 18563 | 0.011* |
| | *gm-PFC* (ln) | 0.089 | -0.014 ▪ 0.188 | 18563 | 0.077 |
| | ***Effect of gm PFC on Brain mass*** | | | | |
| | Intercept | 3.5325 | 2.720 ▪ 3.959 | 18563 | <0.001*** |
| | *gm-PFC* (ln) | 0.851 | 0.651 ▪ 1.052 | 18563 | <0.001*** |
| | ***Effect of brain mass on STM in same subsample*** | | | | |
| | Intercept | 0.129 | -0.416 ▪ 0.644 | 18563 | 0.523 |
| | *Brain* (ln) | 0.109 | 0.002 ▪ 0.209 | 18159 | 0.042* |
| ***c. Neocortex (N = 26 species)*** | | | | | |
| | ***Effect of neocortex on STM*** | | | | |
| | Intercept | 0.348 | 0.141 ▪ 0.578 | 18991 | 0.004** |
| | Neocortex (ln) | 0.069 | 0.013 ▪ 0.121 | 18563 | 0.036* |
| | ***Effect of neocortex on Brain mass*** | | | | |
| | Intercept | 0.779 | 0.459 ▪1.103 | 19522 | <0.001*** |
| | Neocortex (ln) | 0.916 | 0.840▪ 0.987 | 19074 | <0.001*** |
| | ***Effect of brain mass on STM*** | | | | |
| | Intercept | 0.257 | 0.033 ▪0.487 | 18563 | 0.021* |
| | Brain mass (ln) | 0.084 | 0.032 ▪ 0.137 | 18154 | 0.012* |
| | ***Effect of neocortex on STM (N=10)*** | | | | |
| | Intercept | 0.231 | 0.062▪ 0.392 | 18563 | 0.014 |
| | Neocortex (ln) | 0.103 | 0.060▪ 0.144 | 19446 | 0.001*** |
| ***d. Cerebellum (N = 25 species)*** | | | | | |
| | ***Effect of cerebellum on STM*** | | | | |
| | Intercept | 0.445 | 0.320 ▪ 0.579 | 18563 | <0.001*** |
| | Cerebellum (ln) | 0.0813 | 0.037 ▪ 0.125 | 18563 | 0.004** |
| | ***Effect of cerebellum on Brain mass*** | | | | |
| | Intercept | 2.350 | 2.023 ▪ 2.686 | 18563 | <0.001*** |
| | Cerebellum (ln) | 0.864 | 0.762 ▪ 0.963 | 18563 | <0.001*** |
| | ***Effect of brain mass on STM*** | | | | |
| | Intercept | 0.260 | 0.034 ▪ 0.501 | 18167 | 0.024* |
| | Brain mass (ln) | 0.085 | 0.030 ▪ 0.135 | 18113 | 0.012* |
| | ***Effect of cerebellum on STM (N=10)*** | | | | |
| | Intercept | 0.389 | 0.298▪ 0.485 | 18563 | <0.001*** |
| | Cerebellum (ln) | 0.109 | 0.072▪ 0.143 | 18563 | 0.001*** |

**Table S2.**

Results of the MCMCglmm for the effects of brain size on the performance in the short-term memory test data of ManyPrimates et al. (2022), at three different delays (0, 15 and 30 seconds), using either the full sample (**a**), and the reduced sample (**b**) in which only species with at least 10 individuals tested were included.

| | | Post. mean | lower ▪ upper 95% CI | Effective sample size | pMCMC |
|---|---|---|---|---|---|
| ***Short delay (0s)*** | | | | | |
| | ***a. Using total sample (N=41 species)*** | | | | |
| | Intercept | 0.399 | 0.014 ▪ 0.831 | 16866 | 0.035* |
| | Brain mass (ln) | 0.057 | -0.036 ▪ 0.139 | 17304 | 0.211 |
| | ***b. Using only species with at least 10 individuals tested (N= 11 species)*** | | | | |
| | Intercept | 0.263 | 0.039 ▪ 0.485 | 18563 | 0.030 |
| | Brain mass (ln) | 0.111 | 0.059 ▪ 0.162 | 18563 | 0.001** |
| ***Medium delay (15s)*** | | | | | |
| | ***a. Using total sample (N=41 species)*** | | | | |
| | Intercept | 0.160 | -0.076 ▪ 0.389 | 18563 | 0.153 |
| | Brain mass (ln) | 0.093 | 0.040 ▪ 0.148 | 18563 | 0.003** |
| | ***b. Using only species with at least 10 individuals tested (N= 11 species)*** | | | | |
| | Intercept | 0.077 | -0.219 ▪ 0.375 | 18563 | 0.565 |
| | Brain mass (ln) | 0.116 | 0.045 ▪ 0.184 | 18563 | 0.008** |
| ***Long delay (30s)*** | | | | | |
| | ***a. Using total sample (N=41 species)*** | | | | |
| | Intercept | 0.223 | -0.050 ▪ 0.530 | 18563 | 0.095 |
| | Brain mass (ln) | 0.065 | -0.005 ▪ 0.127 | 17789 | 0.078 |
| | ***b. Using only species with at least 10 individuals tested (N= 11 species)*** | | | | |
| | Intercept | 0.097 | -0.214 ▪ 0.399 | 18563 | 0.472 |
| | Brain mass (ln) | 0.108 | 0.041 ▪ 0.180 | 18563 | 0.009** |

**Table S3.**

Results of the MCMCglmm for the effects of the delay on the performance in the short-term memory test of ManyPrimates et al. (2022), replicating the finding of that study.

| | Post. mean | lower ▪ upper 95% CI | Effective sample size | pMCMC |
|---|---|---|---|---|
| ***STM performance*** | | | | |
| Intercept | 0.601 | 0.342 ▪ 0.851 | 18563 | <0.001 |
| Delay | -0.005 | -0.007 ▪ -0.004 | 18563 | <0.001 |

**Table S4.**

Results of the MCMCglmm for the effects of brain size, delay on the performance, and their interaction, in the short-term memory test of ManyPrimates et al. (2022), replicating the finding of that study. Results shown for the full sample (a) and for the reduced sample (b).

| | Post. mean | lower ▪ upper 95% CI | Effective sample size | pMCMC |
|---|---|---|---|---|
| ***a. STM performance in full sample (N=41 species)*** | | | | |
| Intercept | 0.411 | 0.025 ▪ 0.800 | 18563 | 0.035 |
| Brain size | 0.050 | -0.026 ▪ 0.129 | 18116 | 0.211 |
| Delay | -0.005 | -0.012 ▪ 0.002 | 18563 | 0.125 |
| Brain size × Delay | -0.000 | -0.002 ▪ 0.001 | 18563 | 0.922 |
| ***b. STM performance in reduced sample (N= 11 species)*** | | | | |
| intercept | 0.204 | 0.008 ▪ 0.402 | 18140 | 0.042 |
| Brain size | 0.120 | 0.072 ▪ 0.168 | 17923 | <0.001*** |
| Delay | -0.004 | -0.013 ▪ 0.004 | 19431 | 0.290 |

| | | | | |
|---|---|---|---|---|
| Brain size x Delay | -0.000 | -0.003 ▪ 0.002 | 18563 | 0.700 |

**Table S5**.
Results of MCMCglmm analyses for the effect of brain size and each of the 11 possible selective variables considered by ManyPrimates as independent variables on the average performance on the short-term memory test, including only species with at least 10 subjects (n=11 species). Note that none of the possible selective variables affected the statistical effect of brain size or reached significant values.

| **variable** | **Post. mean** | lower ▪ upper 95% CI | **Effect sample size** | **pMCMC** |
|---|---|---|---|---|
| ***Color vision*** | | | | |
| (Intercept) | 0.148 | -0.078 ▪ 0.391 | 18563 | 0.162 |
| Brain | 0.114 | 0.053 ▪ 0.177 | 18563 | **0.006**** |
| Color vision polymorphic | -0.017 | -0.141 ▪ 0.091 | 18563 | 0.745 |
| Color vision trichromatic | -0.022 | -0.240 ▪ 0.194 | 18563 | 0.801 |
| ***Vocal repertoire*** | | | | |
| (Intercept) | 0.142 | -0.030 ▪ 0.321 | 19770 | 0.083 |
| Brain | 0.110 | 0.055 ▪ 0.165 | 19535 | **0.003**** |
| Vocal repertoire | 0.001 | -0.006 ▪ 0.007 | 18105 | 0.855 |
| ***Group size*** | | | | |
| (Intercept) | 0.128 | -0.044 ▪ 0.290 | 19400 | 0.097 |
| Brain | 0.126 | 0.078 ▪ 0.176 | 19615 | **0.001***** |
| Group size | -0.002 | -0.007 ▪ 0.003 | 18563 | 0.360 |
| ***Home range*** | | | | |
| (Intercept) | 0.145 | -0.031 ▪ 0.315 | 18563 | 0.076 |
| Brain | 0.112 | 0.072 ▪ 0.156 | 19097 | **0.001***** |
| Home range | 0.000 | -0.000 ▪ 0.000 | 19003 | 0.850 |
| ***Day journey length*** | | | | |
| (Intercept) | 0.153 | -0.024 ▪ 0.330 | 18563 | 0.074 |
| Brain | 0.104 | 0.045 ▪ 0.162 | 18177 | **0.005**** |
| Day journey length | 0.000 | -0.000 ▪ 0.000 | 17577 | 0.655 |
| ***Resting time*** | | | | |
| (Intercept) | 0.106 | -0.100 ▪ 0.301 | 18563 | 0.231 |
| Brain | 0.115 | 0.076 ▪ 0.155 | 18563 | **0.001***** |
| Resting time | 0.001 | -0.002 ▪ 0.003 | 18563 | 0.450 |
| ***Feeding budget*** | | | | |
| (Intercept) | 0.141 | -0.042 ▪ 0.331 | 18563 | 0.104 |
| Brain | 0.113 | 0.072 ▪ 0.154 | 18563 | **0.001***** |
| Feeding budget | 0.000 | -0.003 ▪ 0.003 | 18114 | 0.991 |
| ***Dietary breadth*** | | | | |
| (Intercept) | 0.094 | -0.034 ▪ 0.238 | 18563 | 0.127 |
| Brain | 0.105 | 0.074 ▪ 0.136 | 19407 | **0.000***** |
| Dietary breadth | 0.018 | -0.003 ▪ 0.038 | 18563 | 0.074 |
| ***Diet diversity*** | | | | |
| (Intercept) | 0.100 | -0.195 ▪ 0.361 | 18563 | 0.310 |
| Brain | 0.115 | 0.049 ▪ 0.193 | 17602 | **0.012*** |
| Frugivore | 0.031 | -0.039 ▪ 0.103 | 18563 | 0.283 |
| Gummivore | 0.105 | -0.063 ▪ 0.287 | 18883 | 0.144 |
| Insectivore | -0.014 | -0.149 ▪ 0.122 | 19453 | 0.727 |
| Omnivore | 0.058 | -0.080 ▪ 0.195 | 18141 | 0.271 |
| ***Frugivory*** | | | | |
| (Intercept) | 0.133 | -0.039 ▪ 0.309 | 19131 | 0.100 |
| Brain | 0.111 | 0.069 ▪ 0.150 | 19299 | **0.001***** |
| Percent frugivory | 0.000 | -0.001 ▪ 0.002 | 18563 | 0.692 |

| | | | | |
|---|---|---|---|---|
| ***Terrestriality*** | | | | |
| (Intercept) | 0.137 | -0.036 ▪ 0.318 | 18563 | 0.094 |
| Brain | 0.115 | 0.071 ▪ 0.159 | 18563 | **0.002**** |
| Terrestriality | -0.008 | -0.100 ▪ 0.077 | 16709 | 0.834 |

Note that repeating the analysis with ln-transformed home range size did not affect this outcome.

**Figure S1.** The relationship between a species' average performance in the short-term memory test and (ln) cerebellum size (**a**) and (ln) neocortex size (**b**) in the reduced sample of at least 10 individuals per species, based on MCMCglmm analyses. See Table S1 for the statistics.

(a) (b)

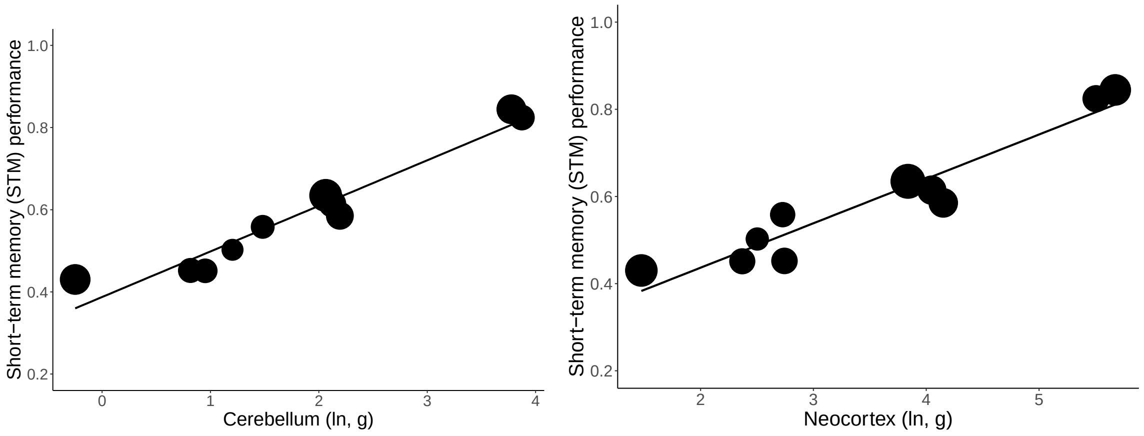
